# Model-Based Reinforcement Learning for Ultrasound-Driven Autonomous Microrobots

**DOI:** 10.1101/2024.09.28.615576

**Authors:** Mahmoud Medany, Lorenzo Piglia, Liam Achenbach, S. Karthik Mukkavilli, Daniel Ahmed

**Affiliations:** Acoustic Robotics Systems Lab, Institute of Robotics and Intelligent Systems, Department of Mechanical and Process Engineering, ETH Zurich, Switzerland; IBM Research - Europe, AI and Accelerated Discovery, Switzerland

## Abstract

AI has catalyzed transformative advancements across multiple sectors, from medical diagnostics to autonomous vehicles, enhancing precision and efficiency. As it ventures into microrobotics, AI offer innovative solutions to the formidable challenge of controlling and manipulating microrobots, which typically operate within imprecise, remotely actuated systems—a task often too complex for human operators. We implement state-of-the-art model-based reinforcement learning for autonomous control of an ultrasound-driven microrobot learning from recurrent imagined environments. Our non-invasive, AI-controlled microrobot offers precise propulsion, which efficiently learns from images in data-scarce environments. Transitioning from a pre-trained simulation environment, we achieve sample-efficient collision avoidance and channel navigation, reaching a 90% success rate in target navigation across various channels within an hour of fine-tuning. Moreover, our model initially successfully generalized in 50% of tasks in new environments, improving to over 90% with 30 minutes of further training. Furthermore, we have showcased real-time manipulation of microrobots within complex vasculatures and across stationary and physiological flows, underscoring AI’s potential to revolutionize microrobotics in biomedical applications, potentially transforming medical procedures.

## Introduction

Artificial intelligence (AI) has significantly advanced capabilities across a variety of fields, including diagnostics^1,2^, fluid mechanics^3,4^, medical imaging^5–7^, and segmentation^8,9^. AI has also played a crucial role in the development of sophisticated drone technology^10^ and autonomous vehicles^11^, reshaping our approach to transportation and surveillance. AI’s integration into microrobotics presents a new frontier^12–14^, introducing distinct challenges in control and functionality. As this integration evolves, successfully navigating these challenges will be key to unlocking the transformative potential of microrobots.

Microrobotic manipulation offers groundbreaking possibilities, from microassembly to surgical microrobotics tasks, improving localized drug delivery systems through precise manipulation within complex vasculatures under physiological conditions^15,16^. However, these applications demand high precision in microrobot control, complicated by their small size and complex dynamics—including system non-linearities, environmental interactions, and external forces like magnetic fields or ultrasound waves that unpredictably affect the motion of microrobots. The manipulation of microrobots, therefore, presents unique challenges surpassing those of traditional robotics. While autonomous vehicles navigate reliably with established technologies like LiDAR and GPS^11^, replicating these sensory capabilities at a microscale is exceptionally challenging. Microrobots often rely on imaging modalities, as most sensors are difficult to scale down or integrate. Furthermore, unlike autonomous vehicles, powered by simpler motor controls, microrobots rely on less precise, wirelessly actuated systems such as light^17–19^, electric^20^,chemical^21–23^, magnetism^24–32^, and ultrasound^33–39^, complicating control further under the influence of external forces.

The deployment of AI in microrobotics also confronts issues like overfitting, vulnerability to errors in new scenarios, and long, often impractical training periods for model optimization. Reinforcement learning (RL)^40^ has proven a powerful tool, enabling robots to learn and adapt directly from their environment. While RL offers potential to surpass human capabilities in tasks like object manipulation and strategic gameplay^41,42^, its reliance on extensive interaction for stable training introduces unique challenges for microrobots, where experimental conditions are highly uncontrolled and variable.

Recent studies have explored a variety of AI-driven microrobots. Researchers have engineering light-driven artificial microswimmers equipped with Q-learning algorithms, which enable them to navigate and learn within noisy environments, effectively overcoming the challenges posed by Brownian motion^43^. Further advancements include the use of deep learning to autonomously steer magnetic swarms, allowing these microrobots to adjust their trajectories through channels of varying dimensions by modifying their shapes^44^. Further innovations have seen the manipulation of magnetic microrobots in 3D using Proximal Policy Optimization (PPO), following training in simulation environments^45^. Spiral magnetic microrobots have also been controlled using deep RL^46^.

Ultrasound-driven microrobots^47–51^, emerging as an exciting non-invasive alternative, are capable of generating tunable propulsive forces, enabling deep navigation into tissues. Nevertheless, achieving precise control and manipulation of these microrobots continues to pose significant challenges, as multiple piezotransducers need to be controlled with millisecond resolution for effective steering—a task often too complex for human operators. Recently, ultrasound microrobots have utilized Q-learning to navigate in a free environment^52^, while adaptive methods have been utilized to control individual particles^53,54^. However, the training time required for manipulation using these algorithms increases exponentially with complexity, a challenge these algorithms find difficult to manage efficiently. Despite this progress, the capabilities for autonomous obstacle avoidance and counter-current navigation remain largely underexplored. Given the incomplete understanding of ultrasound microrobot motion, model-based reinforcement learning (MBRL) is emerging as a promising strategy to navigate them through complex environments with high precision.

This study employs the Dreamer V3 MBRL^55^ algorithm to autonomously control an ultrasound microrobot. Our approach integrates an in-house Python code for imaging and dynamic frequency adjustment of piezoelectric transducers in an artificial vascular channel setup as shown in **Fig 1a**, which controls the ultrasound waves. Additionally, the code interfaces with an electronic circuit designed for rapid switching between transducers, a feature critical for precise navigational steering. Advanced image processing techniques are used to segment images, detect swarms, and track them in real time. This approach frames the microrobots control as a RL task that enhances their performance over time, as shown in **Fig 1b** and **c**. To minimize the need for extensive physical experimentation, we implemented Dreamer V3 to train within an imagined model **Fig 1d** with an actor-critic RL architecture^56^. Although model convergence remains a challenge, taking up to ten days, we developed a Pygame-based simulation environment, as shown in **Fig 1e**, to accelerate the learning of essential navigation skills such as path planning and obstacle avoidance. This knowledge is then applied to enhance system adaptability in real experimental settings, where the system adapts in approximately two hours. Within the simulation environment, we evaluated the MBRL’s performance against state-of-the-art model-free RL (MFRL)^57^ algorithms, demonstrating MBRL’s excellence in managing complex channel navigations where MFRL falls short. As previously discussed in research^55^, the MBRL ability to imagine and simulate future actions as shown in **Fig 1f** and **g** significantly reduces the training time exponentially.

**Fig. 1.**
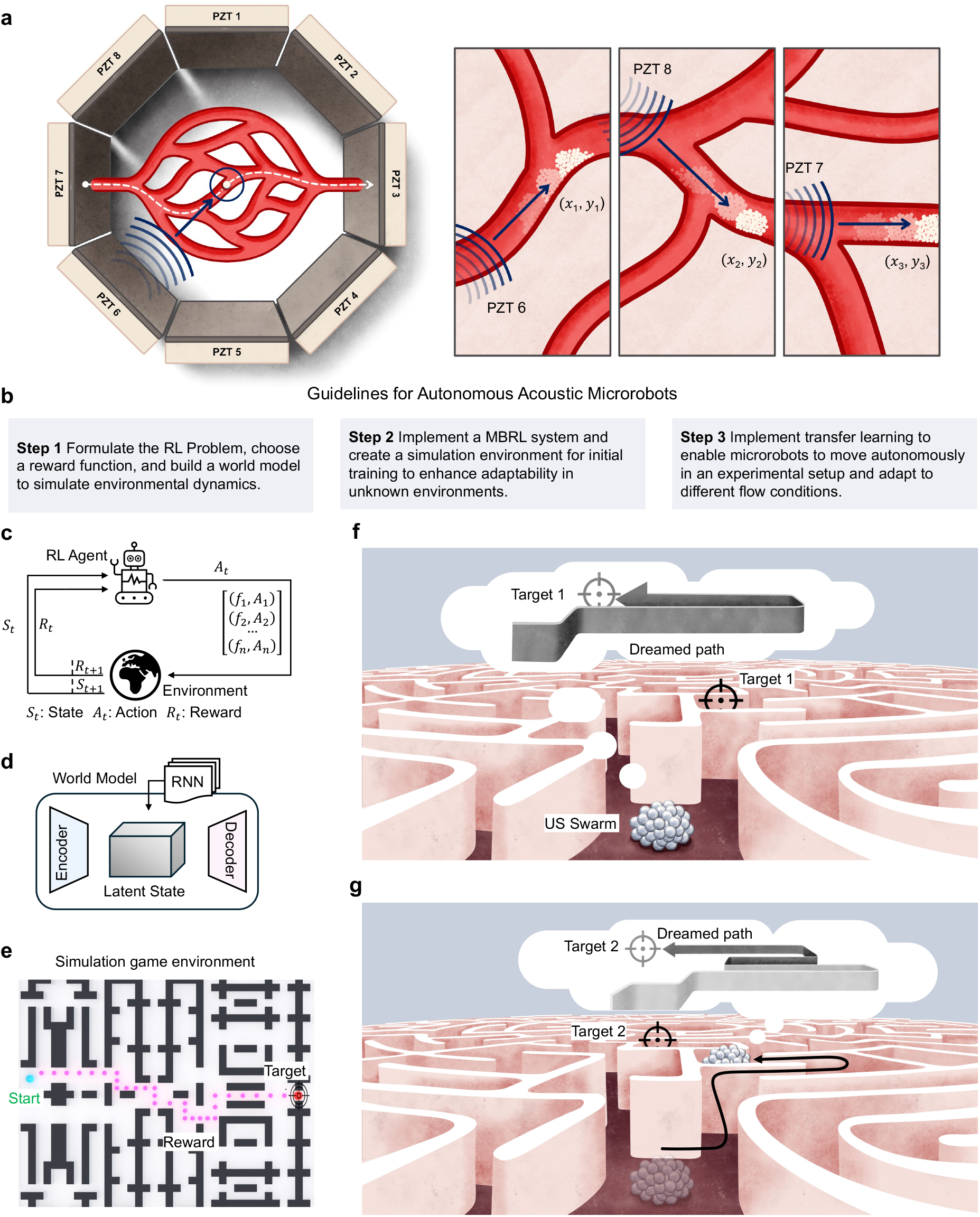
Concept of an autonomous ultrasound-driven microrobot. **a**. Schematic of the experimental setup, showcasing an artificial vascular channel with eight piezoelectric transducers (PZTs) in an octagonal configuration (left image). A schematic illustrates the microrobot’s behavior under ultrasound activation and details methods for its manipulation (right image). **b**. Guidelines for Manipulating Microrobots. **c**. High-level formulation of the Reinforcement Learning (RL) problem, where the environment is our artificial channel and the RL agent is the microrobot with *S*_*t*_, actuated through ultrasound frequency and amplitude to achieve a reward *R*_*t*_ by optimal actions, *A*_*t*_. **d**. A world model encoder-decoder structure has been built to simulate and imagine future states in the environment. **e**. A simulated game environment has been designed to pre-train the microrobots, thereby reducing the convergence time during experimental training. **f**. The microrobot envisions multiple potential paths towards a target in a dreamed environment, allowing it to ‘dream’ and train on various possible scenarios simultaneously. a recurrent ‘dream’ in the latent space **g**. Microrobot agent executes the optimal action, successfully reaches Target 1, and proceeds towards Target 2 using a newly imagined path.

To address potential overfitting from training on a single channel, we have developed a general model trained across diverse channel environments, including vascular structures, racetracks, and mazes. This model consistently delivers 90% accuracy across all trained channels. When tested on a new, previously unseen channel, the model initially achieves a 50% success rate, which remarkably increases to 90% after only 30 minutes of additional training. We further conducted steering tests with microrobots in stationary flow through multiple channels containing obstacles to thoroughly assess and demonstrate the model’s effectiveness. To optimize the model for dynamic flow conditions, we revised the reward function, enabling the microrobots to adhere to walls and navigate within the no-slip boundary to significantly reduce drag. Moreover, by adjusting the amplitude to counteract drag when moving against the flow and reducing it when moving with the flow, the microrobot is capable of advancing against the flow at speeds reaching up to ∼10 mm/s. These results highlight the potential of MBRL in advancing microrobotics for biomedical applications.

## Results

To investigate the control of microrobots through MBRL, we designed an experimental setup that includes an artificial vascular channel, encircled by eight piezoelectric transducers (PZTs) set in an octagonal layout. To control the precise activation and deactivation of the eight transducers, we engineered a custom-built electronic circuit, achieving millisecond switching, integrated with a function generator. The artificial vascular channels were fabricated from transparent polydimethylsiloxane (PDMS) using standard mold replication and soft lithography techniques^58^. The entire assembly is mounted on an inverted microscope. Experimental results—including images and videos—were captured using a DSLR camera recording between 6-18 frames per second, with image acquisition and processing handled by in-house python code.

Our microrobots are produced through the self-organization of commercially available, biocompatible microbubbles, each 2−5 µm in diameter, in an ultrasound field. These microbubbles are introduced into the channel via a controlled liquid pump. In their quiescent state, absent of ultrasound stimulation, the microbubbles remain randomly dispersed within the water solution. However, when subjected to an acoustic field, the microbubbles begin to scatter the sound waves. This scattering, coupled with the synchronized phase oscillation of adjacent microbubbles, triggers their self-assembly^8^. For more details about the experimental setup, please refer to Supplementary Note 1.

To elucidate the steering mechanism of our microrobots positioned near piezoelectric transducer 6 (PZT_6_), we begin by activating PZT_6_. This generates a pressure gradient between the microrobot and the transducer, driving the microrobot from the higher-pressure area towards the lower pressure along the wave’s propagation path, perpendicular to Transducer 6. Our experimental results have shown that microbubbles are highly responsive to ultrasound; even at an excitation voltage of 4 V_PP_, velocities in the range of millimeters per second were achieved in stationary flow. Upon the microrobot’s arrival at the designated checkpoint (*x*_1_, *y*_1_), we proceed by activating the second piezo transducer, PZT_8_, while simultaneously deactivating PZT_6_. This action redirects the steering of the microrobots to align normal to PZT_8_ to continue guiding the microrobot along the intended trajectory (*x*_2_, *y*_2_). We then activate a third piezo transducer, PZT_7_, to steer the microrobot along trajectory (*x*_3_, *y*_3_), concurrently deactivating PZT_8_. This sequence of precise activations and deactivations among the piezo transducers enables sophisticated control over the microrobots’ trajectory, facilitating complex navigational maneuvers, as illustrated in **Fig. 1**.

Achieving precise navigational control over the ultrasound-driven microrobot presents significant challenges to a human operator, primarily due to the need for fast (milliseconds) and precise adjustments in the amplitude and frequency of the ultrasound signal, as well as the activation of various PZT elements. These adjustments influence the behavior of the microrobots, which is often unpredictable. For example, proximity to a specific transducer necessitates a lower voltage to initiate motion, while a microrobot that is farther away requires a higher voltage to be mobilized. Similarly, smaller microrobots require less power, while larger ones need more. Furthermore, while the velocity of microrobots tends to scale linearly with voltage amplitude^52^, their response to frequency adjustments exhibits complex characteristics including Gaussian distributions and multiple peaks, indicating a varied response across different operational frequencies. Adding to the complexity, individual PZT transducers exhibit varied frequency outputs, and the operational frequency ranges of the microrobots differ. Additionally, when microrobots become immobilized, a large amplitude ultrasound spike is applied to effectively dislodge them. This intricate interplay of control parameters underscores the complexity of managing microrobot movements within this advanced experimental framework. The vast variability in voltage, frequency, and switching between PZTs introduces a substantial action space, complicating the control process and necessitating a significant amount of experimental data to effectively navigate this expanded action space.

This vast action space necessitates the implementation of a MBRL strategy. Our approach begins by feeding the MBRL model an image of the vascular channel following any PZT activation. This image acts as feedback for our MBRL model, enabling it to assess the current state of the microrobot within the experiment. We then apply advanced image processing techniques to detect and track microrobot’s movements. The imaging process begins with an inverted microscope, which transmits live images to our processing pipeline (**Fig. 2a**). We segment the initial image into channels and obstacles using the segment anything model (SAM)^59^, chosen for its ability to accurately differentiate complex visual elements. Following segmentation, we refine and clean the image with a morphological closing operation and adaptive thresholding to identify microrobots, which appear black under the microscope. We then apply detection and tracking algorithms to identify the agent (microrobot), calculate the microrobot’s center, and plot a bounding box around it, and initialize the channel and spatial reliability tracker (CSRT) algorithm, selected for its robust tracking capabilities in dynamic and cluttered environments. When tracking is lost, the system quickly re-detects and initiates tracking, minimizing computation and enhancing real-time feedback. In the processed image, microrobots are marked in blue, and the target location in red. Positive rewards are assigned if the microrobot progresses toward the target, while movement away incurs a negative penalty.

**Fig. 2.**
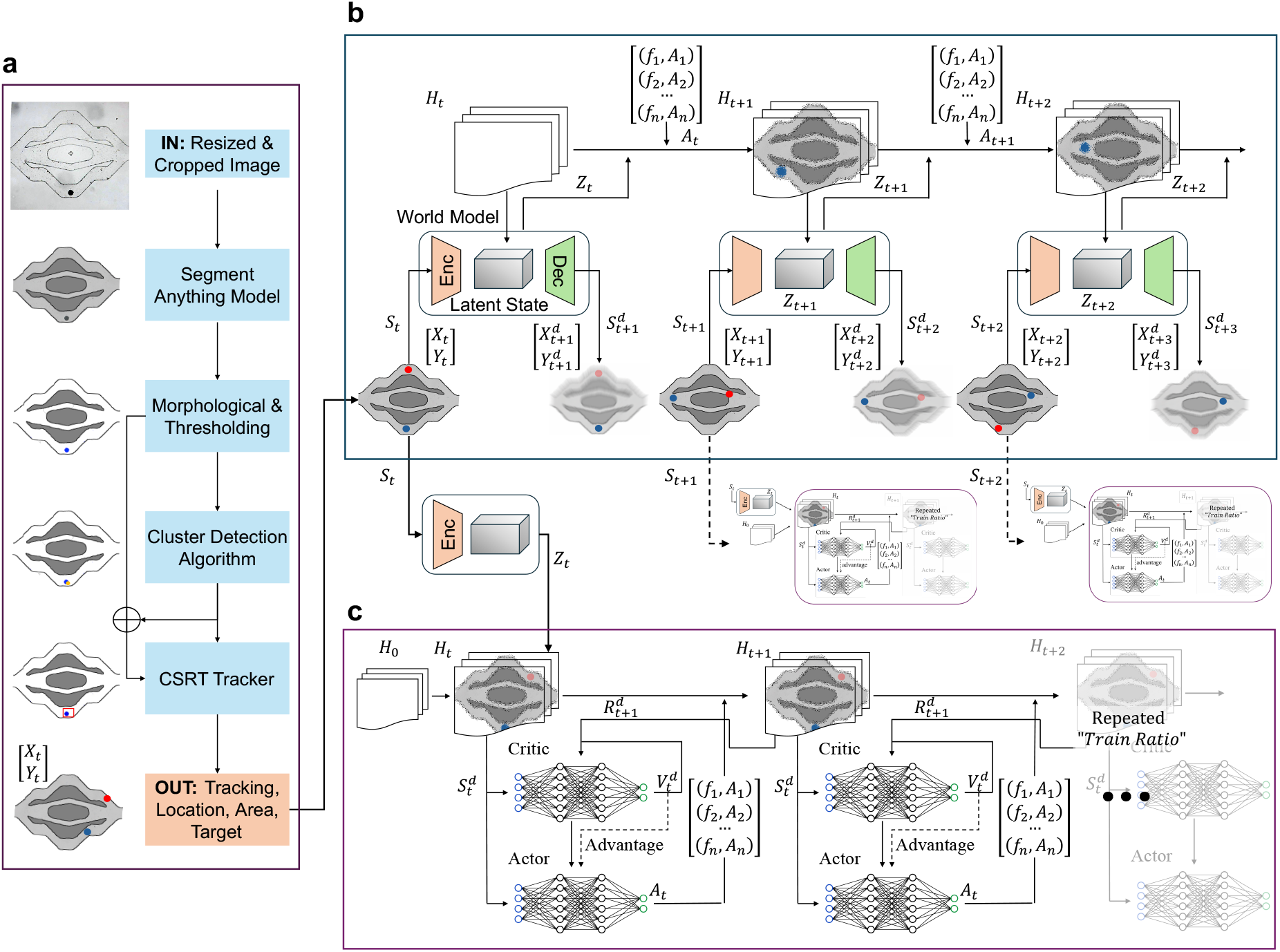
Model-based RL structure for autonomous microrobot. **a**. The imaging pipeline processes channel images to track microrobot positions. It begins by resizing and cropping the images, followed by segmentation using SAM. Morphological operations and thresholding refine the results. A cluster detection algorithm then identifies relevant clusters, and a CSRT tracker monitors the microrobot’s position, providing data on location, area, and target information to the World Model. **b**. The World Model learns to encode the visual output from the imaging pipeline into a latent representation. It employs an encoder-decoder architecture to reconstruct the images, effectively capturing the latent state (*z*_*t*_) from the input state (*s*_*t*_). World model simulates future states (*s*_*t*+1_, *s*_*t*+2_, …) and actions (*a*_*t*_, *a*_*t*+1_, …) in the latent space, facilitating the prediction of microrobot movements. **c**. The actor-critic network is iteratively trained within this loop to optimize the policy, operating in the ‘dreamed’ environment simulated by the World Model. This approach reduces the reliance on real-time interactions with the experimental environment.

### Model-based RL in Microrobotics

In our experimental setup, we employed MBRL to address challenges associated with microrobot manipulation in a complex microvasculature environment. The control problem of the microrobot was formulated using an RL framework, where the state space is defined by image data and the action space as either discrete or continuous variations in frequency, amplitude, and the number of PZT activations. Each action involves adjustments in the amplitude and frequency of a single, specific PZT at a time. Rewards are calibrated based on the microrobot’s efficacy in reaching a target. Details are explained in Methods.

A drawback of MFRL is its dependence on extensive training and interaction with theenvironment. This approach is impractical for our purposes due to the need for a large volume of experimental data to accurately represent the size, quantity, and interactions between microrobots and channel walls in ultrasound settings. Additionally, the slow frame rates required for collecting sufficient images, and the manual interventions needed to adjust microrobot sizes. In contrast, MBRL facilitate more efficient policy learning with fewer samples, making it better suited for complex experimental conditions

To further augment the MBRL setup, we chose the Dreamer V3^55^ algorithm, known for its adept handling of high-dimensional state spaces and complex dynamical systems. This algorithm integrates three primary components: world model learning, envisioning possible future scenarios (**Fig. 2b**), and applying RL to these scenarios (**Fig. 2c**). Together, these elements construct a latent (hidden) model of the environment that predicts or simulates future trajectories, assisting in training decision-making networks with these predictions (see also **Methods** for more details). This approach is particularly valuable as it reduces the reliance on extensive real-world data, which is beneficial in scenarios like microscale operations where data acquisition is challenging. This MBRL setup not only enhances the precision of microrobot control but also optimizes learning efficiency, making it a quintessential tool for advancing robotic interventions in microvascular environments.

### Simulation Environment

We utilized MBRL to train our model using experimental data over a period of six hours. Despite this effort, the model’s performance did not improve, which we attributed to uncertainties surrounding the optimal reward function for this setup. To reduce the need for repeated physical experiments while testing various reward functions, we developed a Pygame^60^-based simulation environment to model the microrobot behaviour. Pygame is a versatile library used to create interactive game environments, which we utilized to simulate dynamic and interactive environments for our microrobots. This environment focuses primarily on local path planning and obstacle avoidance, intentionally omitting the complex dynamics of microrobots, such as PZTs resonance and microbubble size, which are intended for exploration in future real-world experimental setups.

The simulation environment renders as a 64 × 64 RGB image, structured within a ‘gymnasium.spaces.Box’^61^ with dimensions of (64, 64, 3). In this image, obstacles are distinctly marked in dark grey, channels in white, target points in red, and the agent’s position in blue. The agent is depicted with a circular red dot, designed to mimic the microbubble clusters observed in real experimental scenarios. We evaluated the performance of MBRL (Dreamer V3) against PPO^57^, a state-of-the-art MFRL algorithm. Our findings highlight that MBRL exhibited superior efficiency and adaptability within our specific environment, as shown in **Fig. 3**.

**Fig. 3.**
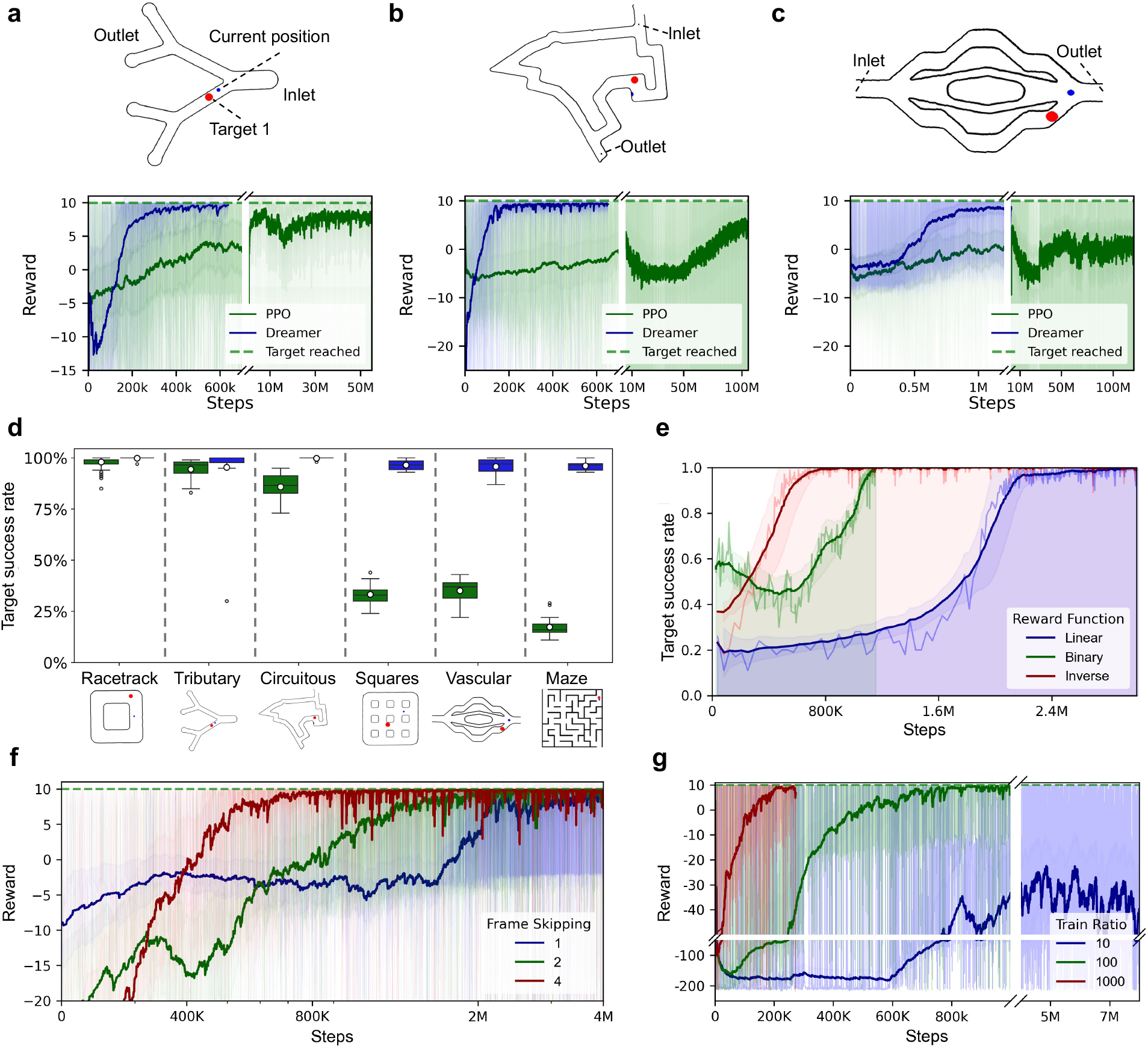
Performance analysis of microrobot navigation in various environments using reinforcement learning algorithms. Reward trajectories for: **a**. Multi-output Tributary channel, **b**. Circuitous channel, **c**. Vascular channels with Dreamer V3 (blue) outperforming state-of-the-art PPO (green) algorithms across simulation steps. **d**. Comparison of PPO and Dreamer algorithms in reaching targets across different channel types: Racetrack, tributary, SPA, Squares, Vascular, and Maze. **e**. Impact of different reward functions on the rate of target achievement. **f**. Effects of frame skipping on performance, presented in a logarithmic plot. **g**. Influence of training ratios on reward dynamics, highlighting consistent performance across various ratios in simulated environments.

**Fig. 3a** demonstrates that in a simple multi-output tributary channel, both algorithms reached convergence; however, our model converged ∼25 times faster than PPO. As the complexity of the task increased In **Fig. 3b**, set against a circuitous racetrack, PPO required up to 100 million steps to converge, whereas MBRL achieved convergence in just 600,000 steps. Further increasing the complexity of the channel, as demonstrated in the vascular example in **Fig. 3c**, MBRL converged after 1 million steps, while PPO failed to converge even after 100 million steps. Overall, our MBRL approach consistently surpassed PPO in performance across a variety of channels, including the complex maze presented in **Fig. 3d**. We experimented with various reward functions, including binary, inverse, and logarithmic, and tested different coefficients for these functions. **Fig. 3e** shows a comparison of these reward functions and highlights the motivation for using an inverse reward function to achieve fastest convergence. This simulated setting enables us to refine and iterate on our reward functions and control strategies efficiently, without the continuous need for live experimental adjustments. This approach streamlines the development process, facilitating more precise and effective advancements in microrobot control. The reward function is designed to incentivize the microrobot to efficiently reach designated target points while navigating around obstacles, taking into account various shapes and layouts of the channels. Formally, the reward *R* at time step *t* is defined by the following criteria:

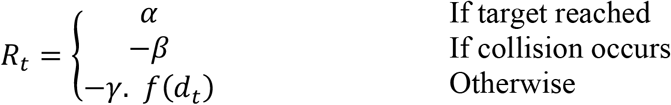

where *α, β*, and *γ* are coefficients that weight the importance of each component in the reward function. The term *d*_*t*_ denotes the Euclidean distance to the target point at time step *t*, and 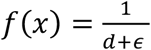 is a real, monotonic function that translates the distance into a penalty (or reward) value, where *ε* is a small positive constant used to avoid division by zero. This function is specifically chosen to inversely relate the reward to the distance, thereby encouraging the microrobot to minimize this distance (see also Supplementary Note 2).

After extensive experimentation, we identified the optimal settings for our system: α = 10, *β* = 2, γ = 0.1. Our simulation results validated that MBRL effectively learns advanced navigation tactics through interactions within the environment, thereby capable of mastering complex navigational strategies in intricate settings such as vascular systems, mazes, and racetracks.

### Implementation of Action Space for MBRL

To effectively train the microrobots, we began by assessing potential combinations of actuator frequencies and amplitudes through experiments, which revealed a large action space with 4 frequency inputs, 4 amplitude inputs, and 4 transduction activation units—resulting in 64 distinct combinations. After thorough analysis of experimental data, we developed an amplitude predictor designed to tailor ultrasound field intensity to the size of the microrobot: smaller swarms require lower amplitudes to avoid overshooting, while larger swarms need higher amplitudes. This concept can be attributed to the scaling of the primary radiation force with the volume of the microrobot. This concept allowed us to refine our approach by eliminating 4 amplitude options, thereby reducing our action space to 16 precise settings. The optimization allows for precise control over microrobot velocity and overshooting based on the size of the microrobot. We further optimized performance by selecting operational frequencies between 2.7 MHz and 2.9 MHz, which align closely with the resonant frequencies of the actuators, to enhance system performance and efficiency.

Initial attempts to augment the latent vector with additional data beyond the processed images, such as the positional coordinates of the microrobots and their target locations. However, this approach did not yield the expected improvements and instead hindered both training and task execution. To enhance the efficiency of our MBRL model, we implemented frame skipping ^41,62^ as a strategy to reduce the computational load and accelerate training time without compromising performance. This method significantly reduces the number of frames processed by the MBRL agent, allowing the model to focus on noticeable changes and minimize the risk of overfitting. Additionally, we implemented max pooling (selecting the maximum pixel value across the skipped frames) across the last two frames to decrease temporal resolution while maintaining essential dependencies between skipped frames. This adjustment greatly enhanced training stability, resulting in smoother convergence, faster learning rates, and improved overall task performance. For our experiments, after testing different frame skipping rates, we opted for 4-frame skipping to achieve faster convergence, as shown in **Fig. 3f**. We also investigated a critical parameter known as the “train ratio,” which denotes the number of steps trained in “imagination,” i.e., within the world model—relative to each step in the “real” environment. This approach capitalizes on the world model to simulate numerous hypothetical scenarios, thus reducing the need for extensive real-world interactions. The key advantage of a higher train ratio is its potential to significantly enhance the efficiency of the learning process. It enables the agent to learn from imagined experiences, which are both quicker and less costly in terms of simulation than real-world interactions. Ideally, a higher train ratio reduces the number of required environmental interactions to achieve convergence.

We experimented with various train ratios to assess their impact on learning efficiency and performance. For example, a train ratio of 10:1 means that for every experimental step, the agent performs ten steps in the dreamed environment. This strategy enables the agent to accumulate more experience and optimize its policy without the time and resource constraints associated with real-world training. Conversely, a lower train ratio, such as 1:1, entails that the agent performs an equal number of real and simulated steps, which significantly slow down the learning process but provides more accurate feedback from the real environment.

Our experiments demonstrated that higher train ratios, such as 1000:1, dramatically reduced the number of interactions with the real environment required to achieve convergence, as illustrated in **Fig. 3g**. The results indicated that higher ratios led to faster convergence, whereas lower ratios often fail to reach convergence. To maximize the benefits of high train ratios, we developed a parallel script to run real environment interactions and world model training on separate threads. This results in an adaptive train ratio that dynamically adjusts with the agent’s performance in the real environment.

### Transition of autonomous microrobots from simulation to real-world environment

Pre-trained models have been highly effective in our physical experiments, adeptly handling tasks like path planning and localization. The main task was adjusting these models to control the frequencies and amplitudes needed to direct microrobots in new settings. Deployed in real-world experiments as shown in **Fig 4a,b** these pre-trained models achieved about 70% of our target objectives within 10 days of continuous operation, though the full training extended to 16 days. This extension was due to occasional microrobot failures when unmonitored and issues with managing microrobot sizes; when microrobots became too small, they would overshoot and lose track, prompting the introduction of new microbubbles to rebuild the microrobots. Additionally, our experimental protocol required us to pause every 12 hours to perform experiments on freshly self-assembled microrobots. The variability in the diameters of these new microrobots, which rarely matched, prevented the model from overfitting to any specific microrobot size. This variability ultimately enhanced the model’s ability to generalize, making it more robust and adaptable to different swarm sizes (see also **Supplementary Note 3**).

**Fig. 4.**
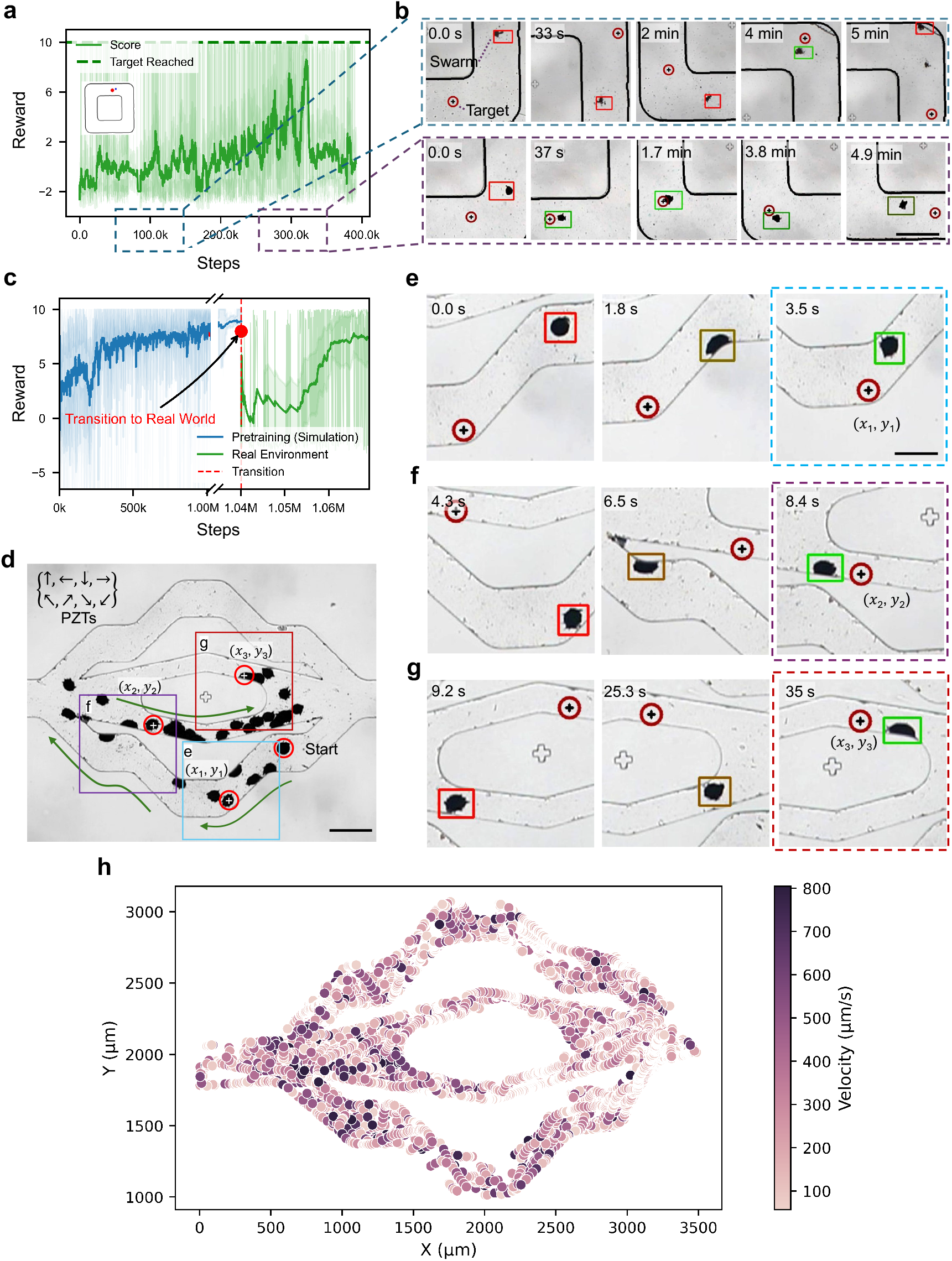
Transition of autonomous microrobots from simulation to real-world environment. **a**. A plot of MBRL demonstrates training of microrobots with 32 actions in an experimental racetrack square channel. The *y*-axis indicates the reward function, with positive values signalling successful navigation towards the target and negative values indicating crashes. The *x*-axis displays the number of training steps. The reward drops significantly at around 300k steps due to overfitting to a specific corner of the channel. **b**. (Top) Image sequences illustrate the experimental training of microrobots at 100k steps of graph **a**. At this early stage, microrobots frequently fail and reach the target only 20% of the time. (Bottom) The image sequence shows that by 350k training steps, the microrobots demonstrate improved efficiency, reaching the target faster (in seconds) 75% of the time. A red box marks the microrobot when it is far from the target, while green indicates proximity to the target. The target position is denoted by a circle with a black plus symbol. **c**. A plot of RL performance illustrates the transition of a pre-trained model from simulation to the real-world environment. The score rapidly increases during the simulation phase (blue), drops after the transition to experiments (marked by a red vertical dashed line), and then increases as the model adapts to the real-world environment (green). **d**. A micrograph with superimposed images of the microrobots’ navigation steps within an artificial vasculature, with the start and target points marked by red hollow circles. Arrows in the top left corner indicate all possible movement directions of the microrobots. **e, f, g**. Image sequence demonstrating a microrobot navigating an artificial vasculature, starting from the initial position and sequentially reaching three predefined targets: (*x*_1_, *y*_1_), (*x*_2_, *y*_2_) and (*x*_3_, *y*_3_). **h**. The heatmap depicts the relationship between speed and position within an artificial vasculature channel. The color gradient indicates the speed at each position, with darker colors indicating higher speeds and lighter shades denoting slower velocities.

The model, when trained solely on experimental data, exhibited a tendency to overfit, causing the microrobots to remain within certain sections of the channel. Forcing the microrobots to navigate to different areas of the channel resulted in decreased performance and failure to reach the designated targets. This behavior underscores the need for further model adaptation to minimize overfitting. It also highlights the value of incorporating pre-training in a simulated environment, which can significantly reduce training time and improve the model’s overall adaptability.

### Mapping Continuous Actions within a Discrete Framework

While pre-training on discrete actions initially offered benefits, it was constrained by a limited action space—choosing only four frequencies, with varied responses across PZTs. Realizing the importance of determining the optimal frequency for each PZT for effective microrobot navigation at each channel point, we transitioned to continuous actions. However, the existing pre-trained models on discrete actions proved unsuitable. The variability of each PZT and their interactions with the PDMS channels introduced significant challenges in accurately simulating the necessary physical interactions for training on continuous actions.

To overcome these challenges and facilitate precise adjustments in frequency and amplitude, we implemented a continuous action space. This space includes frequency values ranging from 2.7 to 2.9 MHz and amplitudes from 4 to 14 V_PP_, selections informed by manual control experiments. Additionally, we incorporated the rapidly exploring random tree star (RRT*) algorithm for path planning, which ensures optimal navigation paths. By activating a PZT opposite to the desired direction of movement, we were able to focus exclusively on learning the nuances of continuous actions for frequency and amplitude.

To ensure that the agent consistently followed the designated paths, we refined the reward function, triggering a reset and initiating a new episode whenever significant deviations from the designed path occurred. During training, we observed a saturation of the reward function, primarily due to repeated frequency adjustments. These adjustments often caused the microrobots to overshoot their targets, resulting in erratic movements. Furthermore, the model frequently opted for higher amplitudes in an attempt to accelerate target acquisition. While this approach initially seemed advantageous, this strategy typically increased navigation instability, i.e., further deviating from the target point. The frequent fluctuations in both frequency and amplitude compromised precise control, making it challenging to accurately adhere to the designated paths.

The complex and dynamic environment, coupled with the system’s nonlinearities and intricate action space, presented significant challenges in achieving consistent, stable performance from the model. Despite these challenges, integrating continuous action with path planning, we manage to navigate the microrobots, though not perfectly, as demonstrated in **Supplementary Note 4**. Additionally, a significant issue emerged when paths ended; the RRT* algorithm’s delay in computing new paths often resulted in microrobots being swept away by the flow in the vasculature channel.

In response, we incorporated a sweeping action around the resonant frequency of the PZT into each discrete action using our programmable function generator set to 1 ms steps. This strategy leverages the resonant frequency characteristics of the PZTs to ensure that microrobots consistently operate at or near their optimal frequencies. By utilizing pre-trained environments and optimized paths, we enhanced performance and reduced experimental time (**Fig. 4c**). The implementation of sweeping actions resulted in smoother transitions and more stable movements, addressing the overshooting issues observed during the continuous training phase, as shown in **Fig. 4d,e,f** and **g**. This refined approach represents a significant advancement in microrobot navigation, enabling more precise and effective control. **Fig. 4h** presents a heatmap illustrating the relationship between speed and position within an artificial vasculature channel. The color gradient indicates speed, with darker shades representing higher velocities and lighter shades indicating slower ones.

### Dynamic Adaptation of MBRL to Complex and Variable Environments

We have demonstrated that once a model is trained to navigate a particular environment, it is capable of adapting to diverse environments through a fine-tuning process. Initially, transitioning from a simple to a complex maze required approximately 100,000 steps, typically spanning an hour (see also **Fig. S6**). To reduce the adaptation timeframe and prevent overfitting to any single training scenario, we exposed the model to a variety of environments, including various vascular networks, mazes, and racing circuits. **Fig. 5a** illustrates the reward function and the target reach score across ten mixed environments. Subsequently, the model was able to adapt to an entirely unknown environment, such as transitioning to the “multi-output tributary channel,” within about 50,000 steps, or roughly 30 minutes. Moreover, **Fig. 5b** quantitatively illustrates the model’s ability to achieve target objectives across a range of training environments over various training steps. It highlights a progressive increase in the rate at which targets are reached, demonstrating improvements from 25% to 100% as the model accrues more training across environments, from simple constructs like “Empty” and “Quadrant channel” to more complex configurations such as “Maze Medium” and “Maze Hard”. The analysis reveals both the adaptability of the model and its increasing proficiency over time, especially notable in complex and challenging environments. This progression underscores the effectiveness of the model’s learning mechanisms in adapting to diverse and increasingly difficult scenarios, reflecting its potential for robust applications in dynamic settings. However, it is noteworthy that as the complexity of the environments increases, the model’s convergence time tends to lengthen.

**Fig. 5.**
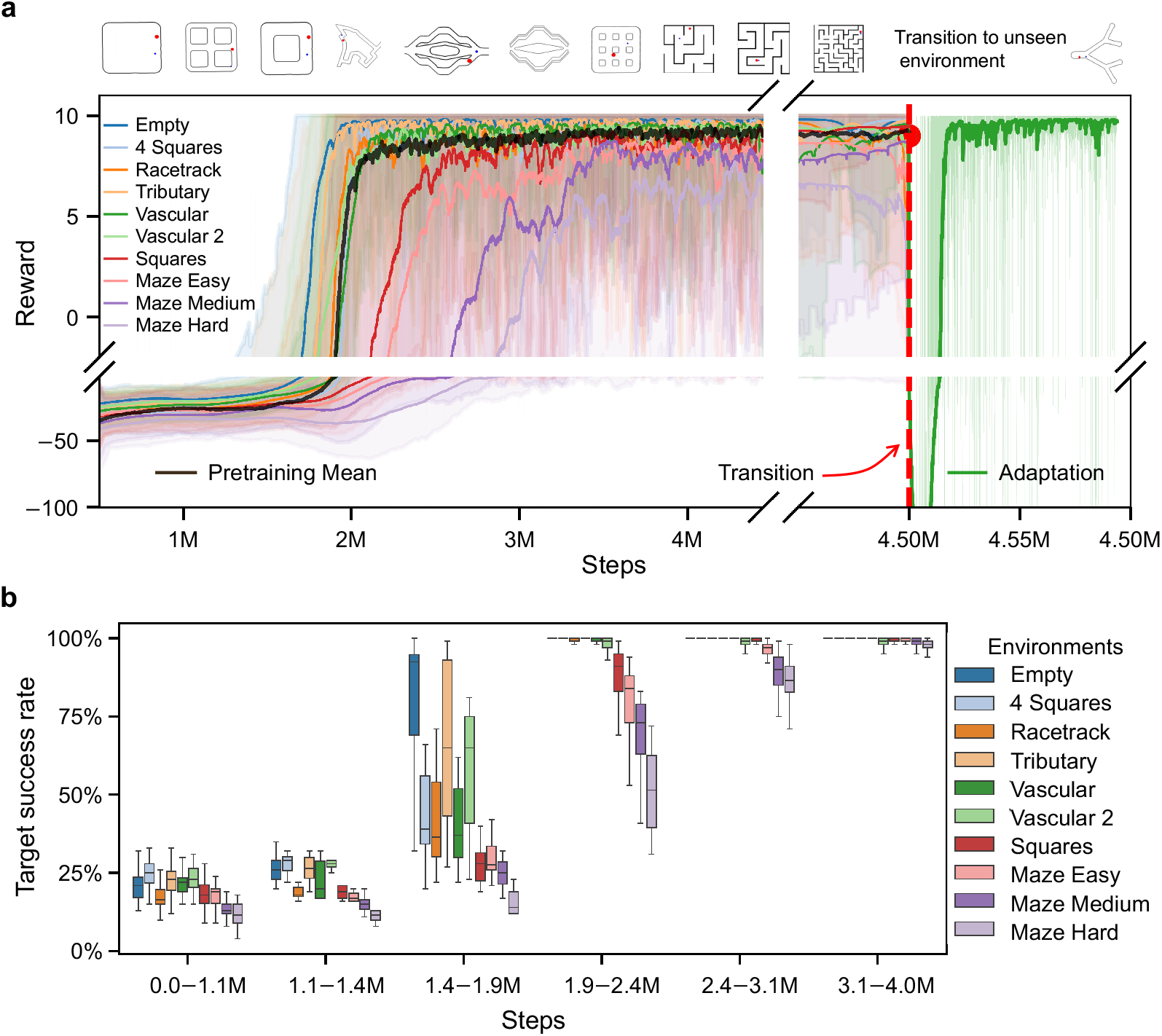
Performance of the generalized world model across various environments. **a**. The plot presents the reward progression over training steps across nine distinct environments, including Empty, 4 Squares, Racetrack, Vascular, and various Maze configurations. Each line on the graph denotes the average reward for each environment, effectively illustrating the model’s learning trajectory. Simpler environments achieve convergence faster, whereas more complex ones require additional time, but all eventually reach convergence. The region beyond 4.5 million steps, demarcated by a dashed vertical red line, marks the phase of pretraining and subsequent adaptation to a new environment. This transition point highlights the shift from pretraining across ten simulation environments to adaptation within a new multi-output tributary channel. The model demonstrates significant improvement over the initial 4.5 million steps of pretraining, followed by rapid adaptation, achieving stable performance within just 50,000 steps (approximately 30 minutes) in the new environment. **b**. The success rate of targets reached across different environments is plotted against training steps. Box plots illustrate the variability and distribution of the MBRL algorithm’s performance in successfully reaching targets. While simpler environments facilitate quicker convergence, our MBRL model consistently attains convergence across all scenarios.

### Autonomous manipulation in physiological flow

Autonomous navigation and manipulation within dynamic flow environments present significant challenges for microrobotics^63–65^. Initially, our models were trained under no-flow conditions within a vascular channel; however, when these models were subsequently applied to flow conditions, they faced considerable difficulties due to the increased drag forces. The flow often flushed away microrobots, necessitating substantial manual effort to restart the training and assembly of the microrobots. Furthermore, the pre-trained model was now less effective due to the bigger domain gap between the simulation and the real environment.

We first adjusted the reward function to impose penalties for microrobots moving into the center of the channel (see also **Supplementary Note 5**), where drag forces are typically strongest as shown in **Fig. 6a**. The adapted reward function *f*(*dt, Xt, A*_*t*_) is defined as follows:

**Fig. 6.**
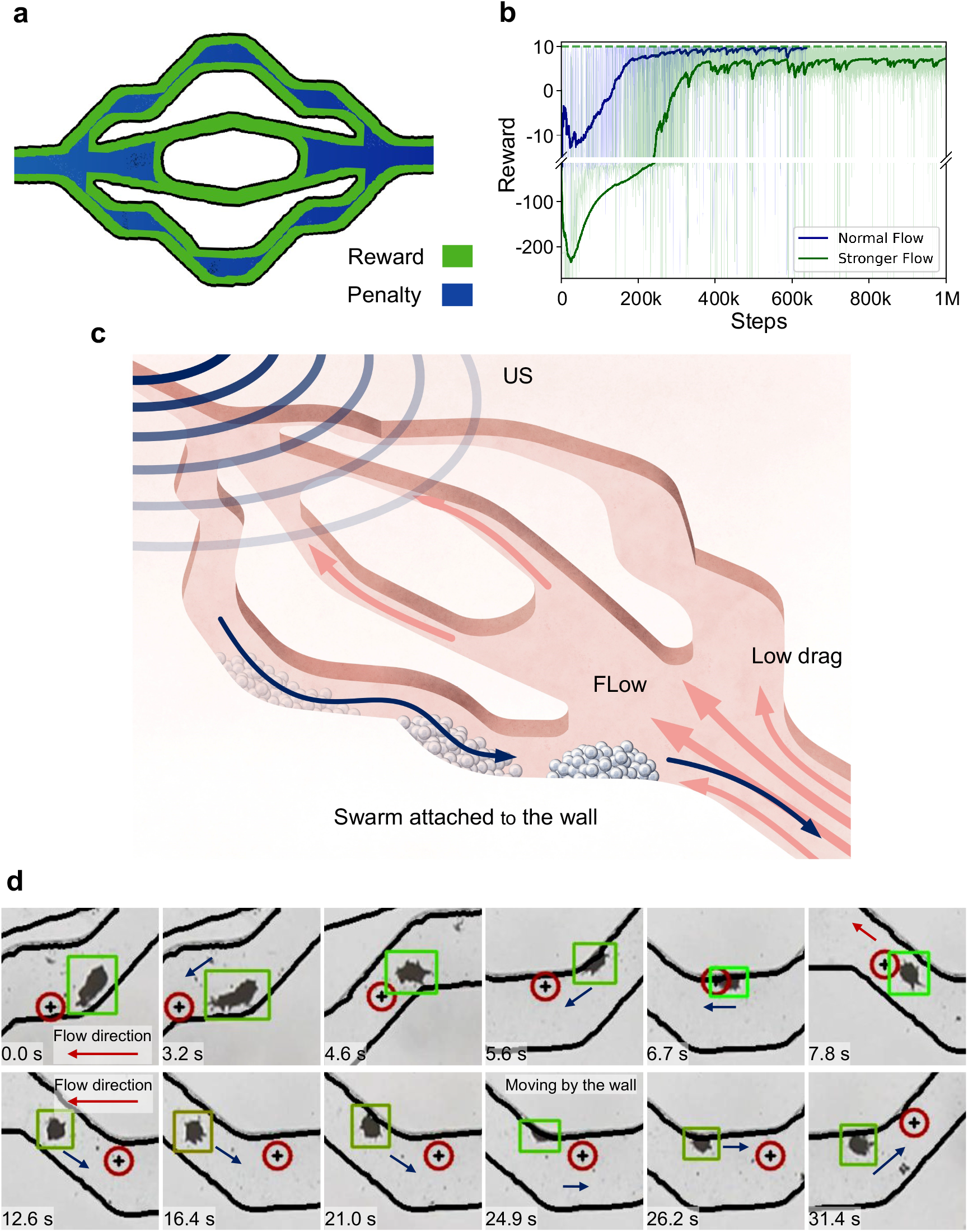
Autonomous microrobot navigation upstream in a flow environment. **a**. Schematic of the reward function adjustment to promote microrobot navigation close to the wall, minimizing drag. **b**. Graph showing reward progression over time for microrobots in normal (blue line) and stronger (green line) flow conditions, highlighting differences in learning and adaptation. In normal flow, rewards steadily improve and stabilize around 200,000 steps. In stronger flow, initial difficulties lead to more negative rewards, but the algorithm shows significant improvement by 400,000 steps. **c**. A schematic illustrates the behavior of the microrobot attached to the wall to avoid drag and move against the flow. **d**. An image sequence shows a microrobot navigating within a microfluidic channel under flow conditions. Initially, the microrobot encounters maximum drag, i.e., when it is at the center of the channel (0.0 s to 6.7 s) before moving towards the wall (7.8 s to 31.4 s), where the drag is minimal. The reduced drag forces at the wall facilitate more stable and controlled navigation. The red arrows indicate the direction of the fluid flow.

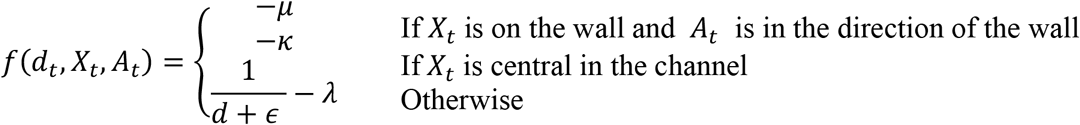

The components of the reward function are initialized as follows: a step penalty *λ* is applied at each step to encourage the microrobot to reach the target quickly. The wall sliding penalty −*µ* is imposed when the microrobot is in contact with a wall and the action taken is in the direction of the wall, allowing for sliding along the wall but discouraging pushing against it. The inverse distance reward 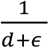 provides a continuous incentive for the microrobot to move closer to the target, with stronger gradients as the distance decreases. Moreover, a centering penalty −*κ* is introduced for being too centrally located in the channel, encouraging the microrobot to stay near the walls where drag forces are lower.

These adjustments incentivize microrobots to navigate closer to the channel walls, where drag forces are significantly reduced due to the no-slip condition. We also incorporated a simulated force in our environment that continuously pushes against the microrobots, quantified in pixel values corresponding to the flow rate we aim to counteract. As shown in **Fig. 6b**, stronger flow requires more steps to achieve convergence. These adjustments allow for realistic simulation of the physical challenges encountered by microrobots in flow conditions. Finally, we refined the physical model of microbubble dynamics, focusing particularly on bubble-wall interactions. Specifically, the secondary Bjerknes force induces microbubble cluster attraction and subsequent adhesion to the channel walls as shown in **Fig .6c**. The cluster then benefits from the “no-slip” condition at the wall, which produces reduced shear forces and facilitates easier movement along the wall.

In addition to environment characteristics, the response of microrobots to acoustic actuation— particularly how they react linearly to changes in voltage and cluster size—necessitated adjustments to the amplitude predictor. Our strategy for navigating environments with flow involves increasing power when moving against the flow to counteract the increased drag and reducing power when moving with the flow to take advantage of the reduced resistance. This differential power strategy ensures more efficient navigation and manipulation within complex flow environments as shown in the image sequence in **Fig 6d**. Finally, in the event we lose sight of the microrobots, we have implemented a rescue function that accesses their last coordinates and attempts to reverse the recent actions to return the microrobots to our field of view.

## Discussion

Ultrasound microrobots, known for their biocompatibility, offer significant advantages for biomedical applications, including minimal invasiveness and the ability to operate in complex environments such as living tissues. Controlling these microrobots, however, remains a bottleneck. In this study, we demonstrate that it is possible to steer microrobots using only ultrasound in complex channels and guide them autonomously against flow in real time by utilizing state-of-the-art MBRL strategies. Moreover, we show that incorporating a simulation environment accelerates this process. Transitioning from a pre-trained simulation environment, we achieve sample-efficient collision avoidance and channel navigation, reaching a 90% success rate in target navigation across various channels within an hour of fine-tuning. Additionally, our model initially generalized successfully in 50% of tasks in new environments, improving to over 90% with 30 minutes of further training. Furthermore, for movement in a flow environment, we adjusted our simulation setup to incorporate fluid dynamics knowledge, exploiting low-drag regions near the wall and the attractive forces between the microrobots and the wall. This enabled real-time movement both against and with the flow, underscoring AI’s potential to revolutionize microrobotics in biomedical applications.

We envision our work being applied across a range of manipulation strategies within microfluidics, enhancing single-cell studies and facilitating manipulation research on small animal models such as C. elegans^33^ and zebrafish embryos^66^. Additionally, our techniques could significantly advance microparticle separation and other precision applications in biotechnology and healthcare. Our approach also holds the potential to drive breakthroughs in microrobotic technology, particularly in minimally to non-invasively invasive surgical procedures. Our approach also promises to catalyze breakthroughs in microrobotic technology, particularly in minimally invasive to non-invasive surgical procedures. This includes the use of ultrasound-driven microrobots, such as shape-morphing^47^ and spiral^48^ designs, as well as those propelled by streaming forces^51^, offering innovative solutions for complex medical interventions. Moreover, the integration of ultrasound with 3D printing^67,68^ could also benefit from our approach, enabling precise particle manipulation for bioprinting applications. For the microrobotics community, our image-based model can also be easily adapted for light^17–19^, chemical^21–23^, electrical^20^, and magnetic^30–32^ actuation systems, following initializing the actuators, enabling autonomous control of diverse microscale objects.

Looking ahead, we anticipate expanding our work into 3D manipulation by incorporating a side camera and refining the imaging pipeline for 3D data acquisition. Integrating medical imaging techniques such as ultrasound and two-photon microscopy will allow us to study microrobot behavior in animal models more effectively^14^. Achieving complex maneuvering and task execution in three-dimensional spaces will require a more sophisticated actuator system. With these technological advancements, we aim to streamline the process for in vivo testing by utilizing state-of-the-art segmentation methods in medical applications^8,9^, starting with animal models like mice^69,70^ and eventually scaling up to larger mammalian models.

## Methods

### Microchannel Fabrication

The microfluidic channels used in the study were produced through standard soft lithography using PDMS. Each device was fabricated using a master mold, lithographically patterned with SU-8 negative photoresist on a 4-inch silicon wafer, which was later placed inside a Petri dish. The thermocurable PDMS prepolymer was prepared by mixing the curing agent with the base at a weight ratio of 10:1. After degassing under vacuum, the prepolymer was cast onto the mold. PDMS was cross-linked by thermal curing for 2 hours at 85°C. The PDMS was poured into a mold and then cut and peeled from the channel mold. A 0.75 mm punch was used to punch the inlet and outlet ports. The ports were created by mounting the puncher at an angle of 60 degrees, which prevented fluid from entering the channel at an angle that may cause a break in the plasma treatment bond between the layers of PDMS, resulting in leakage and malfunctioning of the environment. An additional layer of PDMS was bonded to the PDMS channel by plasma treatment for 1 minute at 85°C for 2 hours. A piezo transducer was attached to the PDMS channel wall orthogonal to the aneurysm cavity. The channel flow was circulated using a pulsatile or continuous flow pump through tubes attached to the inlet and outlet. To avoid the impedance mismatch between the other side of the channel wall and the air, the entire system was placed in a water container.

### RL Implementation

We formalized the problem as a Markov Decision Process (MDP), which includes the state space, action space, reward function, and transition dynamics. The state, action, reward triplet (*S*_*t*_, *A*_*t*_, *R*_*t*_) and the transition dynamics (*T*) enable the RL agent to learn optimal policies for microrobot control through continuous interaction with the environment. The state space *S*_*t*_ incorporates visual information captured by cameras, including the current position extracted from the images 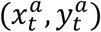 coordinates, and the target location 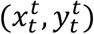 coordinates, representing the spatial location of the desired target position that the microrobot aims to reach represented at time *t* in this equation.

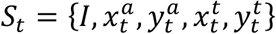

where *I* encapsulates the processed camera feed at time *t*. (CNNs) are employed to extract meaningful features from the images, such as the size, shape, and interactions of the microrobots: *I* = *CNN*(*Image*_*t*_)

The action space *A*_*t*_ defines the set of all possible actions the control system can execute at any given time. In our settings, these actions pertain to the settings of the PZTs: where *f* is the frequency of the US traveling wave, and *A* is the amplitude representing the peak-to-peak voltage and n is number of transducers.

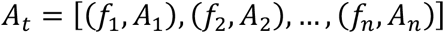

The transition dynamics *T*(*S*_*t*_, *A*_*t*_) describe how the state of the system changes in response to an action. This function is unknown to the RL algorithm and must be inferred through interaction with the environment. In our settings, the transition dynamics represent the physical changes in the system state resulting from the activated PZT. It is expressed as:

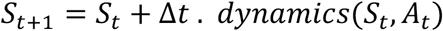

where Δ*t* is the timestep, and *dynamics*(*S*_*t*_, *A*_*t*_) is a function modeling the physics of microrobot motion under ultrasound stimulation, extracted from the differences in the images (state).

### World Model Learning

The world model processes the state *S*_*t*_ into a latent state *Z*_*t*_ using an encoder-decoder architecture. This model predicts future latent states and rewards based on the current latent state and actions. It continuously trains on new samples (*S*_*t*_, *A*_*t*_, *R*_*t*_). The key components include:

▪ Encoder-Decoder Architecture: This architecture compresses high-dimensional observations into a compact latent space for prediction and control. The encoder *q*_*ϕ*_ maps an observation *o*_*t*_ to a latent state *z*_t_, where *ϕ* is the shared parameter vector between the encoder and all other world model components:

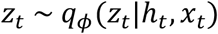 The decoder (*D*) reconstructs the observation from the latent state: *ô*_*t*_ = *D*(*z*_*t*_)
▪ Dynamics Network: This network predicts the future states of the microrobots based on their current state and actions, following the principle of an Recurrent neural network (RNN). It preserves a deterministic state *h*_t_ predicted by the RNN using the previous actions *a*_*t*−1_, *h*_*t*−1_, and the previous embedded state *z*_*t*-1_.

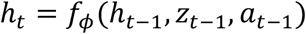
▪ Reward Predictor: This component predicts the rewards associated with different actions, aiding the agent in optimizing its behavior. The reward predictor *R* estimates the reward *r*_*t*_ based on the latent state *z*_*t*_ and action *a*_*t*_.

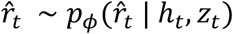

### Latent Imagination and Policy Optimization

The agent generates future trajectories within the latent space and uses these imagined trajectories for policy and value network training. This reduces the need for real-world interactions and allows for more efficient learning. The main steps involved are:

▪ Trajectory Sampling: Generating possible future trajectories by simulating the environment using the transition model (*h*_*t*_ = *f*_ϕ_(*h*_*t*-1_|*z*_*t*-1_, *a*_*t*-1_). The imagined trajectories start at the true model states *s*_*t*_ drawn from the replay buffer of the Agent, then carried in imagination by the transition model. These trajectories are generated much faster than the environment interaction and are controlled by a parameter called “train-ratio”. We developed a multi-threaded approach where the latent model runs continuously on a separate process without a fixed ratio with the real environment interactions.
▪ Trajectory Evaluation: Assessing the quality of each trajectory based on the accumulated rewards predicted by the reward model. The Reward predictor 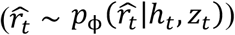 estimates the rewards of each state.
▪ Policy and Value Network Training: The Actor-Critic component is trained to maximize the expected imagined reward 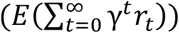 with respect to a specific policy. The evaluated trajectories are used to update the policy and value networks, which dictate the agent’s actions in the real environment.

This training loop leverages the predicted latent states and rewards, significantly enhancing sample efficiency by reducing the dependence on real-world interactions and relying on a very compact latent representation.

#### Algorithm 1

Microrobot MBRL Training

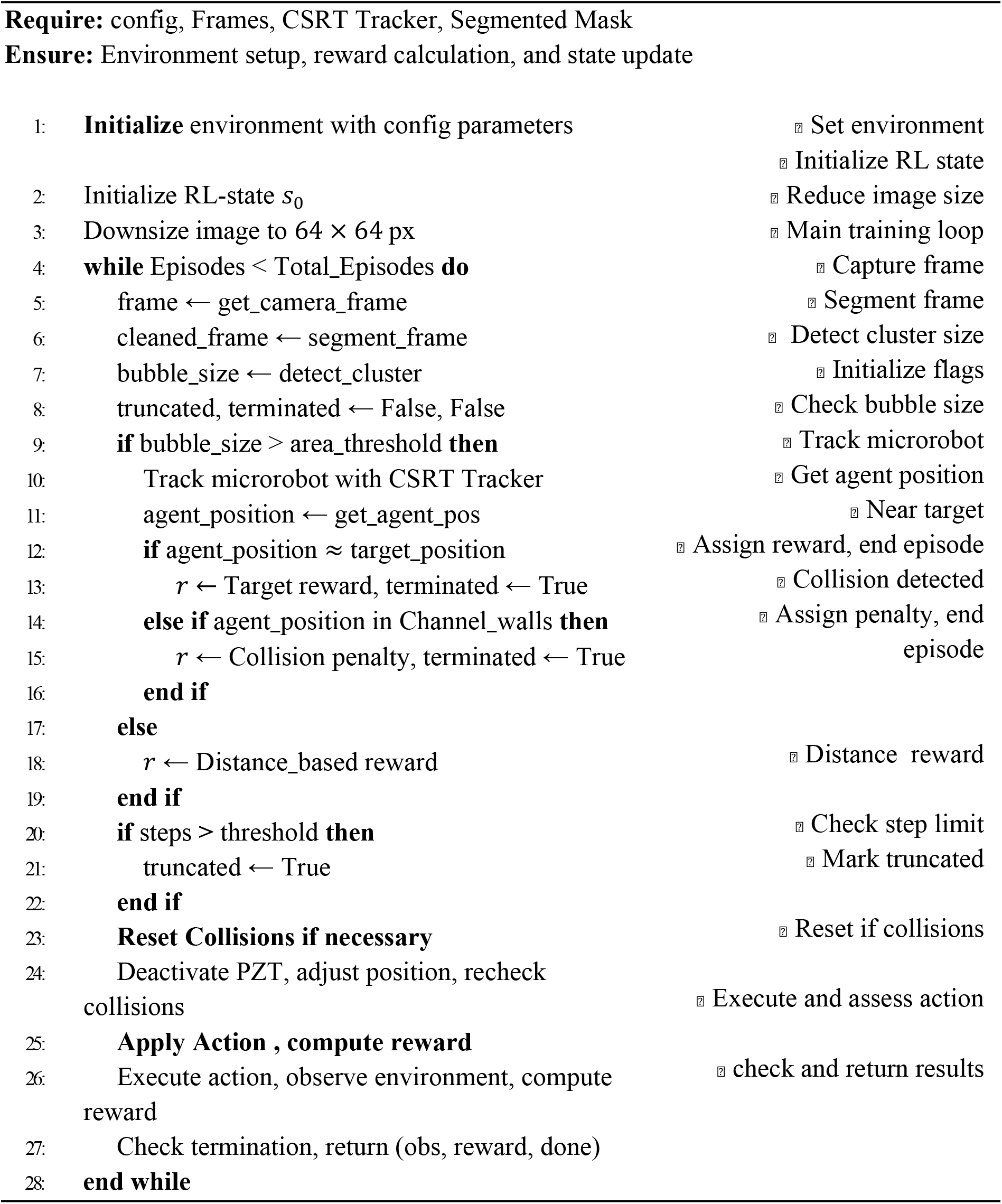

#### Algorithm 2

Microrobot Flow Environment Training and simulation

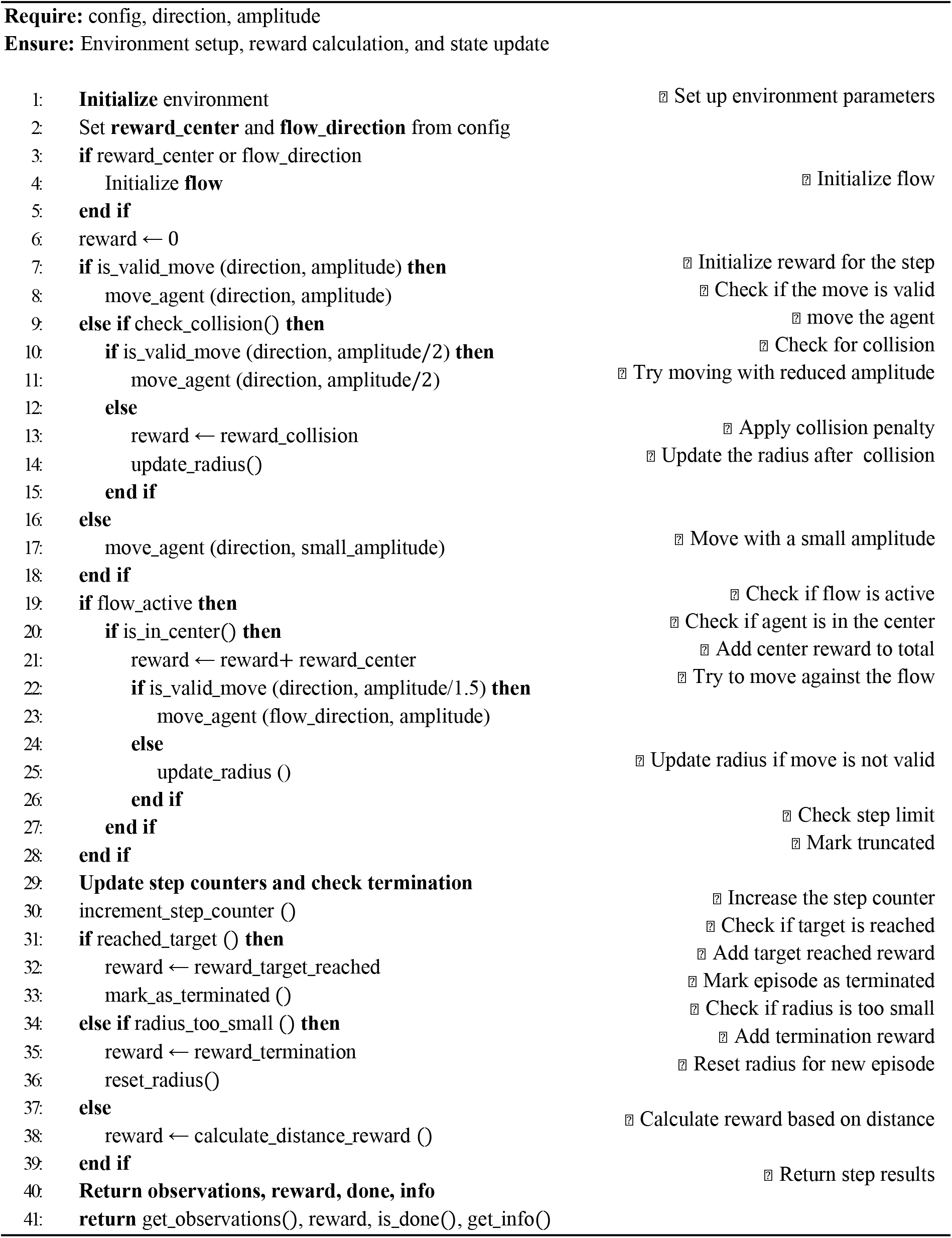

## Contributions

M.M. and D.A. conceived the project. D.A. supervised the project. M.M., L.P, L.A. performed all the experiments and performed data analysis with feedback from K.M and D.A. M.M, L.P., and D.A. contributed to the experimental design, scientific presentation. M.M, L.P., L.A.,K.M. and D.A. wrote and reviewed the manuscript.

## Acknowledgements

This project has received funding from the European Research Council (ERC) under the European Union’s Horizon 2020 Research and Innovation Programme Grant Agreement No. 853309 (SONOBOTS), Swiss National Science Foundation (SNSF) under the SNSF Project funding MINT 2022 grant agreement No. 213058, and ETH Research Grant, grant agreement No. ETH-08 20-1.

