## Supporting Information for "Model-Based Reinforcement Learning for Ultrasound-Driven Autonomous Microrobots"

Contents:

- Supplementary notes S1 to 6
- Extended Data Fig. 1 to 3
- Legends for movies S1 to S6
- SI References

### Note S1. Experimental setup

The experimental setup for the autonomous manipulation of microrobots consists of three main components: the physical setup, imaging pipeline, and control unit.

#### 1. Physical Setup

The physical setup includes a PDMS (polydimethylsiloxane) microfluidic channel, eight piezoelectric transducers (PZTs) as actuators, a function generator, and a microcontroller connected to an electronic circuit containing eight relays responsible for selectively activating and deactivating the PZTs. The function generator (Tektronix AFG3000) is used to generate the sinusoidal waves needed to manipulate the microbubbles. It allows for real-time modulation of acoustic signals, enabling precise control over frequency and amplitude. The microfluidic channels are fabricated using soft lithography techniques, beginning with a master mold patterned lithographically using SU-8 negative photoresist on a silicon wafer, which is then placed inside a Petri dish. The PDMS prepolymer is prepared by mixing the silicon elastomer base and curing agent in a 10:1 weight ratio, degassing the mixture under vacuum to remove air bubbles, pouring it into the mold, and curing it at 85°C for 2 hours. Once cured, the PDMS is peeled off the mold, and inlet and outlet ports are punched using a 1-mm punch. A separate, flat PDMS layer is created similarly using a blank wafer, with both layers cleaned using isopropyl alcohol and subjected to a 15-minute ultrasonic bath in water. After a 1-minute plasma treatment, the layers are aligned and pressed together at 85°C for 2 hours. Eight PZTs, each resonating at 2.8 MHz, are attached to the sides of the PDMS-embedded microfluidic channel using two-component epoxy glue. The microbubbles used are ultrasound imaging contrast agents from Bracco Sonovue[1], which are prepared by injecting saline into a vial containing lyophilized sulfur hexafluoride lipid-type A powder and gently shaking the mixture to create stable microbubbles. These microbubbles have a sulfur hexafluoride gas core surrounded by a phospholipid monolayer shell that prevents coalescence, with sizes ranging from 2 to 9 µm. To prevent standing wave formation and ensure traveling waves, the setup is submerged in water. The microbubbles remain evenly distributed without acoustic actuation, but applying an incident acoustic field causes them to aggregate into swarms due to the secondary Bjerknes force[2,3].

#### 2. Imaging Pipeline

The imaging setup features a Canon EOS 6D Mark II camera mounted on a Leica DMI6000B inverted microscope, capturing live images that are transmitted via HDMI to the processing pipeline. A segmentation model identifies obstacles and navigable spaces within the microfluidic channel, with adaptive intensity thresholding used to detect microbubbles, which appear black under the microscope. The threshold is optimized through iterative trials, depending on camera settings and lighting conditions. Processed images highlight microbubble clusters in blue (0, 0, 255) and plot the current target location in red, matched to the cluster size. The segmentation model is specifically designed to differentiate the background from obstacles in the channel. This process begins with an operator-assisted segmentation model that separates the channel and obstacles (SAM)[4]. This model integrates a robust vision transformer with prompt-based interactions, delivering excellent zero-shot generalization and accurately segmenting new, unseen images without additional training, followed by a morphological closing operation using a (5, 5) kernel over three iterations to refine the segmented mask, which is then applied to the original image. Morphological operations are useful for removing imperfections and smoothing the segmentation results. While different preprocessing methods like adaptive thresholding, Otsu's method, and edge detection techniques (e.g., Canny and Sobel) were explored, the best results were consistently achieved by feeding the raw image directly into the segmentation model. To enhance efficiency, segmentation masks for common channels are cached, reducing the need for repeated segmentation during restarts or resumptions, which significantly cuts down on computational overhead.

**3. Control Unit**

The control unit is a computer equipped with an NVIDIA RTX 4090 GPU with 24 GB of memory, running on Ubuntu. It oversees the MBRL training and manages the entire experimental setup. The control unit integrates data from both the physical setup (including the Arduino and function generator) and the imaging pipeline (capturing video signal from the camera via a capture card). It runs the control algorithms while dynamically adjusting experimental parameters, enabling real-time feedback and precise control. This integration is crucial for ensuring seamless coordination between the various components, allowing for efficient and responsive management of the experimental environment.

### Note S2. Simulation environment and amplitude predictor.

#### Simulation Environment

We developed a simulation environment to model microrobot behaviour, intentionally excluding resonance and bubble size effects. This design decision was based on the simulation's purpose for training rather than replicating microrobots' complex dynamics. The theory was validated in a sample environment resembling the real setup to conserve resources and time. Our primary focus was on local path planning and obstacle avoidance, with the environment configured as a 64 x 64 RGB image, represented by gym.spaces.Box with uint8 values ranging from 0 to 255 and a shape of (64, 64, 3). Obstacles were delineated in black, the channel in white, target points in red, and the agent's position in blue. The agent's circular shape mimicked microbubble clusters observed in real-world scenarios **Fig. S1**. The simulation physics utilized the PyGame framework, providing high efficiency and easy integration with the MBRL environment. This simulator enabled rapid testing of various algorithms[5,6], architectures and parameters, facilitating quick iteration and development of control strategies.

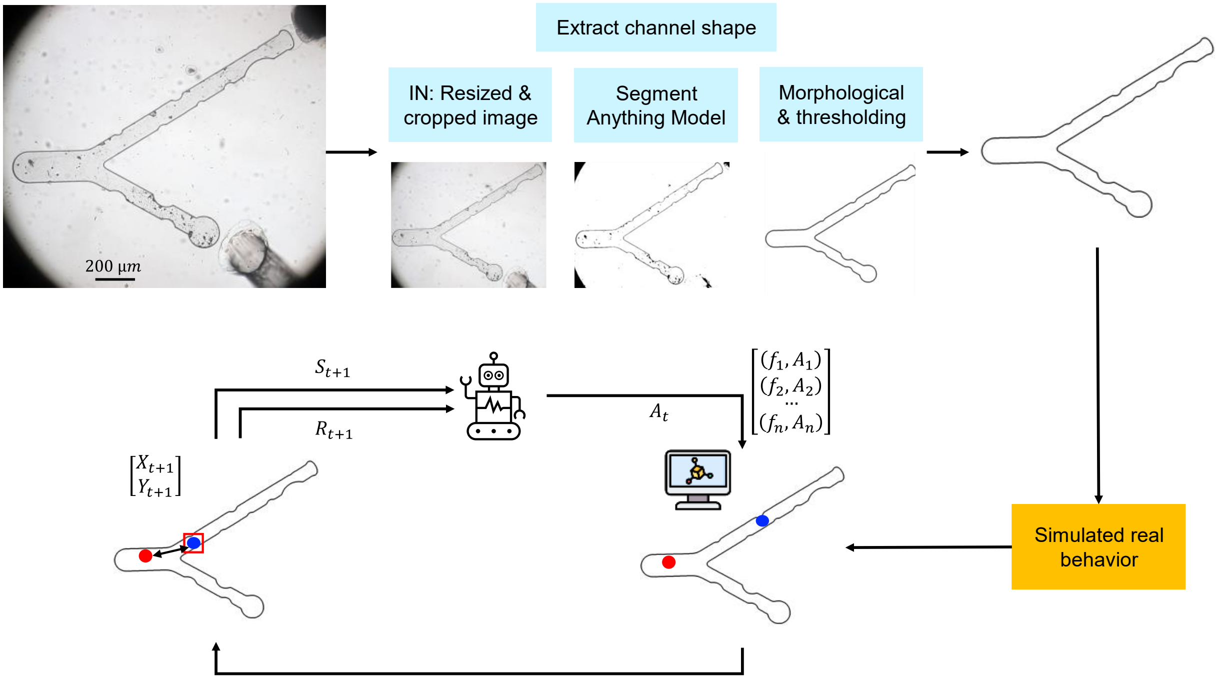


**Fig. S1 | Simulation Environment:** The process begins with extracting the channel shape from real microscope images (top left). The image is resized and cropped, followed by segmentation, morphological processing, and thresholding to identify the channel boundaries (top center). The extracted channel shape is used to create a simulated environment that mimics real-world behavior (middle right). The agent interacts with this simulated environment, receiving state updates and generating actions to navigate the channel (center). The simulator outputs the next state (Xt+1, Yt+1 ) based on the agent's actions (bottom left) and updates the visual representation of the channel (bottom right). This iterative process ensures the simulated behavior aligns with real-world dynamics.

#### 1. Reward Functions

We tested various reward functions to optimize microrobot behavior, balancing penalties for incorrect actions (e.g., wall collisions) and rewards for correct ones (e.g., reaching the target). The general reward function is:

$$R_t = \begin{cases} \alpha & \text{If target reached} \\ -\beta & \text{If collision occurs} \\ -\gamma.\ f(d_t) & \text{Otherwise} \end{cases}$$

where $f(d_t)$ is based on the distance to the target ($d$):

- **Binary Reward Function:** Provides fixed rewards for reaching the target (α = 10) and penalties for crashes ($\beta = 2$), with ( $f(d_t) = 0$). This approach offers clear guidance but may struggle with long-horizon planning due to decaying rewards at larger distances.
- **Linear Distance Reward Function**: Rewards based on linear distance:
  $$f(d) = d + \upsilon \text{ where } (\upsilon)$$
  is a hyperparameter. It provides straightforward guidance but may suffer from diminishing gradients near the target.

- **Inverse Distance Reward Function**: Rewards inversely proportional to the distance:
  $$f(x) = \frac{1}{d + \epsilon}$$
  This function offers continuous feedback, increasing the pull towards the target as the distance decreases.
- **Logarithmic Distance Reward Function**: Uses a logarithmic scale:
  $$f(d) = \log\left(\frac{1}{d + \epsilon}\right)$$
  It offers strong gradients near the target, stabilizing navigation and preventing overshooting.
- **Squared Distance Reward Function**: Employs an inverse quadratic scale:
  $$f(x) = \frac{1}{(d + \epsilon)^2}$$
  This function intensifies the reward as distance decreases, driving precise movement towards the target, but requires careful tuning to ensure stability.

**2. No Collision**

To further refine the reward structure, especially for environments where microrobots slide along walls, we introduced the "No Collision" reward function. This modified function removes harsh penalties for collisions and instead implements small penalties and step rewards to encourage effective navigation while allowing flexibility in handling obstacles.

The reward function is defined as follows:

$$R_t = \begin{cases} \alpha & \text{If target reached} \\ -\gamma.\ f(d_t, X_t, A_t) & \text{Otherwise} \end{cases}$$

$$\text{Where: } f(d_t, X_t, A_t) = \begin{cases} -\mu & \text{If } X_t \text{ is on the wall and } A_t \text{ is in the direction of the wall} \\ \frac{1}{d+\epsilon} - \lambda & \text{Otherwise} \end{cases}$$

- **Positive reward for reaching the target** ($\alpha$): Fixed positive reward for reaching the target.
- **Wall sliding penalty** ($\mu$): Small penalty when the microrobot is in contact with a wall and the action taken is in the direction of the wall, allowing sliding but discouraging pushing against the wall.
- **Inverse Distance Reward** $\left(\frac{1}{d+\epsilon}\right)$: Continuous incentive for the microrobot to move closer to the target, providing stronger gradients as the distance decreases.
- **Small step penalty** ($\lambda$): This constant penalty encourages the microrobot to minimize the number of steps taken to reach the target. It is designed to keep the reward always negative.

These modifications ensure that the microrobot is able to handle collisions gracefully by allowing sliding along walls: it can adapt its shape and path more naturally in response to the fluidic environment.

It is important to note that with this reward function, the cumulative reward for a long episode where the target is not reached can become very negative. This occurs because the reward is always negative until the target is reached, accumulating penalties over time. In contrast, the previous function reset the episode upon collision, incurring only a fixed penalty of $-2$.

**Amplitude Prediction**

We developed an amplitude predictor to reduce the action space by determining the optimal amplitude for the function generator based on the observed area of the micro-robots. The relationship between bubble size and required amplitude was modelled using experimental data.

The amplitude predictor uses a formula correlating the observed area of the microbubbles to the required voltage:

$$A = k \cdot \sqrt{area} + b$$

where A is the amplitude in volts, area represents the area of the microbubbles in pixels, and k and b are empirically determined calibration constants. This approach allows the amplitude to vary from approximately 4 volts for very small bubbles to 18 volts for larger clusters.

Integrating the amplitude predictor reduced the action space from 64 to 16 actions, optimizing control strategy and improving training efficiency without compromising control over the micro-robot's velocity and stability.

### Note S3. Transfer learning from simulation to real environment

#### Pretraining in a racetrack channel

The pre-training phase involves training the reinforcement learning (RL) algorithm within a simulated environment specifically designed to mimic the real-world conditions of microrobot navigation.

- **Enhanced Collision Physics Simulation**: The simulator incorporates a more realistic collision model. In this model, when microrobots contact the channel walls, they flatten slightly, simulating the physical deformation that occurs in real scenarios. This enhancement provides a more accurate representation of the microrobots' behaviour upon impact, improving the reliability of the simulation.
- **Size Randomization**: To mirror real-world variability and prevent the model from overfitting to a specific microrobot size, the simulator randomizes the microrobot size at the beginning of each episode. This ensures that the model is exposed to a range of sizes, promoting generalization across different scenarios.
- **Speed Randomization**: The simulator also introduces variability in the agent's speed at each instant, which is inversely correlated with the frequency of specific actions. This approach encourages the exploration of less frequently used actions, avoiding the model's collapse into a single dominant behaviour. The speed in each direction is sampled as a normal random variable, where the mean is inversely proportional to the soft max of the action frequencies:

$$s_d \sim \mathcal{N}\left(\sigma\left(\frac{1}{f_i}\right)\right), \theta^2)$$

Where:

- $s_d$ is the speed in direction d.
- $f_i$ is the frequency of action iii in the "actions-buffer" (a storage of the most frequent actions).
- $\sigma\ f_i$ is the softmax function applied to the frequency $f_i$ , which converts these frequencies into a probability distribution:

$$\sigma(f_i) = \frac{e^{f_i}}{\sum_{j=1}^{k} e^{f_i}} \quad \text{for } i = 1,2, \dots, k$$

- $K$ is the number of possible directions.
- $\theta^2$ is the variance of the speed, controlling the spread of the normal distribution.

This method works by decreasing the speed in directions that are frequently chosen, thereby encouraging the exploration of less common actions. It helps to avoid overfitting to a specific set of behaviors and increases the entropy (randomness) of the action space, leading to more robust learning across a broader range of conditions. Once the model achieves satisfactory performance in the simulation environment, it is transferred to the real experimental setup. To ensure a smooth transition, the final portion of the replay buffer from the simulation is also transferred.

During the initial adaptation phase, we observe a brief performance degradation due to the domain shift between the simulation and real environments as shown in **Fig.3a**. However, this adaptation period is short, and the model quickly adjusts to the real-world conditions, effectively bridging the domain gap to reach the target as shown in **Fig. 3b**. Importantly, the pre-trained model shows significantly better performance compared to a model trained from scratch in the real environment.

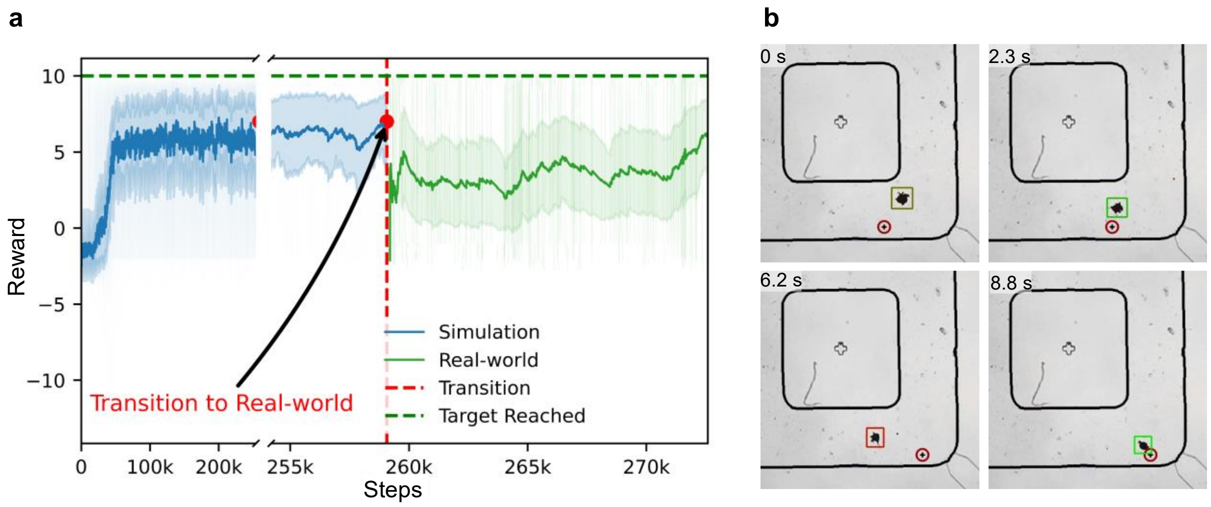


**Fig. S2 | Transfer learning in a racetrack channel. a.** A plot of RL performance illustrates the transition of a pre-trained model from simulation to the real-world environment. The score rapidly increases during the simulation phase (blue), drops after the transition to real-world experiments (marked by a red vertical dashed line), and then rises again as the model adapts to the real-world environment (green). **b.** An image sequence demonstrating a microrobot navigating an artificial racetrack channel, starting from the initial position and sequentially reaching the target. A red box marks the microrobot when it is far from the target, while green indicates proximity to the target. The target position is denoted by a circle with a black plus symbol.

### Note S4. Continuous action space implementation

During training, we observed that frequent adjustments in frequency led to overshooting targets, causing instability in microrobot movements. The model often selected higher amplitudes to reach targets more quickly, which initially appeared beneficial but ultimately increased instability. The dynamic nature of the microrobot's environment, coupled with the nonlinearities and complexity of the action space, presented significant challenges for the model in achieving stable and reliable performance.

Despite these challenges, employing continuous actions and path planning allowed for successful navigation of the microrobots, although not optimally as shown in **Fig. S3**. Frequent oscillations in frequency and amplitude compromised precise control, hindering the microrobot's ability to follow designated paths accurately.

Another key limitation of this approach is the inefficiency of RRT planning: Once a path ended, the RRT* planner required significant time to calculate a new path. Hence, while effective in static conditions, it proved too slow to react to dynamic changes in the environment, such as fluid flow. As a result, while the combination of continuous actions and path planning showed potential, the system's inability to react swiftly in dynamic conditions significantly limited its practical application.

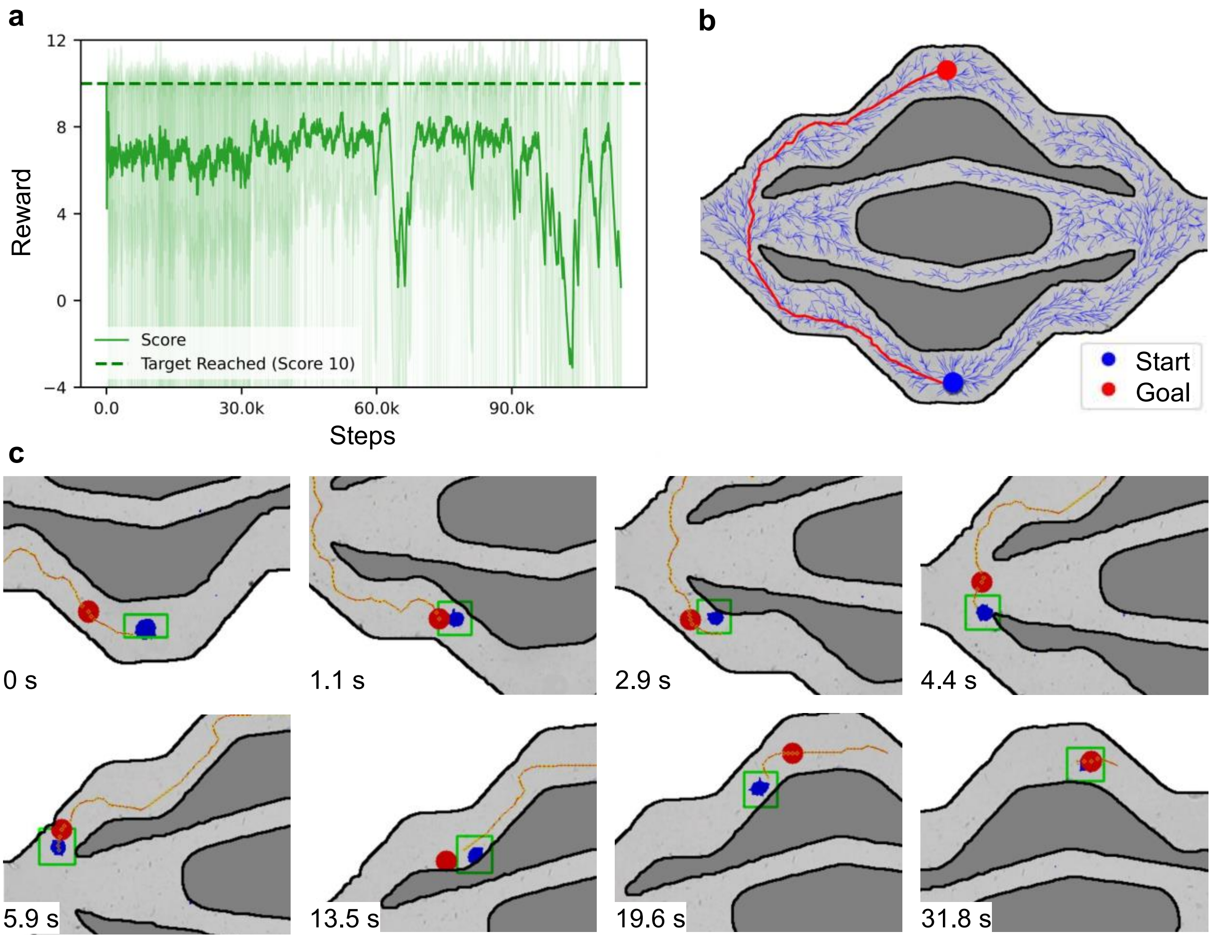


**Fig. S3| Training with Continuous Actions. a.** A plot displaying the relationship between the reward and the steps , illustrating the training progress and performance trends. **b.** RRT* path planning within an artificial vascular channel. **c.** A sequence of images showing the microrobot (blue) following the preplanned path (yellow) to reach updated targets (red). The updated targets and intermediate points are marked along the path to visualize the microrobot's tracking accuracy and performance.

### Note S5. Flow simulation adjustments

Several adjustments have been made to enhance the robustness and adaptability of the microrobots in dynamic flow conditions:

- **Reward Function Adjustments**: To encourage microrobots to avoid areas with high drag forces, we adjusted the reward function to impose penalties for microrobots moving in the centre of the channel. This adaptation incentivizes the microrobots to navigate closer to the channel walls, where the drag forces are significantly reduced due to the no-slip condition.

  The adapted reward function $f(d_t, X_t, A_t)$ is defined as follows

$$f(d_t, X_t, A_t) = \begin{cases} -\mu & \text{If } X_t \text{ is on the wall and } A_t \text{ is in the direction of the wall} \\ -\kappa & \text{If } X_t \text{ is central in the channel} \\ \dfrac{1}{d+\epsilon} - \lambda & \text{Otherwise} \end{cases}$$

  Where we added a Centring Penalty ($\kappa$): A penalty for being too centrally located in the channel, encouraging the microrobot to stay near the walls where drag forces are lower.

- **Physical Model Refinements**: We refined the physical model of microbubble dynamics to better simulate bubble-wall interactions. The second Bjerknes force, which induces attraction and subsequent adhesion of microbubble clusters to the channel walls, was incorporated into the MBRL model. This interaction results in the formation of a "mirrored cluster" effect, allowing microrobots to leverage the reduced shear forces near the walls for easier movement.

- **Simulated Flow Force**: To realistically simulate the physical challenges encountered by microrobots in flow conditions, we introduced a simulated force that continuously pushes against the microrobots. This force is quantified in pixel values corresponding to the flow rate we aim to counteract. By simulating this dynamic force, the microrobots can adapt their strategies to navigate effectively within the flow. Furthermore, this force is stronger the more the bubble is in the center of the channel, behaving according to the Bernoulli equation.

**Note S6. Algorithm for segmentation, reset function and frame skipping**

Algorithm 1: Image Segmentation and Tracking Pipeline

**Require:** Input image from the camera
**Ensure:** Segmented image, Bounding box, and Initialized Tracker

```
Input: Resized & Cropped Image
segmented image ← SegmentAnythingModel(image)
morphed image ← MorphologicalCloseOperation(segmented image, kernel_size)
thresholded_image ← zeros_like(image)
thresholded_image[image ← threshold] ← [255, 0, 0]
cleaned_image ← thresholded_image AND morphed_image
bbox_size ←0
While bbox_size < min_cluster_size do
    bbox, bbox_size ← detect_cluster(cleaned_image)
End While
Tracker ← CSRT_tracker.init(morphed_image, bbox)
Output: Live Tracking, Location, Area
```

Algorithm 2: Collision Reset Function

```
for each step in Collision Reset Steps do                     ▷ Iterate through reset steps
    self.set_piezo_off()                                      ▷ Function generator off
    obs ← self.get_obs()                                      ▷ Iterate through reset steps
    pos ← self.tracker.get_agent_pos(obs)                     ▷ Get microrobot's position
    Count_collisions ← self.check_collisions (obs, pos, radius)
    piezo ←ARGMAX(count_collisions)                           ▷ Select piezo with most collisions
    self.set_piezo_on(piezo)                                  ▷ Activate selected piezo
    if not self.in_collision(pos) then                        ▷ Check if clear of obstacles
        self.set_piezo_off()                                  ▷ Deactivate piezo
        break                                                 ▷ Exit loop
```

Algorithm 3: Frame Skipping Implementation

```
Initialize obs_buffer, total_reward, done
for i in range (skip) do
    obs, reward, done, info ← env.step(action)                ▷Take action
    Store obs[image] in obs_buffer                            ▷Store for max pooling
    Total_reward += reward                                    ▷Sum reward
    if done then
        break                                                 ▷Exit if done
Max_frame ← max(obs_buffer)                                   ▷Max pooling
return max_frame, total_reward, done, info                    ▷Return results
```

### Legends for movies S1 to S6

**Movie S1.** Training process of MBRL in a real microfluidic racetrack channel. The left video shows early training (<100k steps) where the algorithm struggles to reach the target. On the right, after 340k steps, the algorithm shows improved target acquisition. The full process spans 10 days due to real-environment interactions.

**Movie S2.** Transfer learning behavior from a simulation environment to a real experimental environment of the same shape. It shows that the model converges in real experiments in just 3 hours.

**Movie S3.** Continuous action training in a vascular channel. The left side shows RRT* blue tree branches searching for the shortest path, marked in red when found. On the right, the microrobot is marked in blue, with the next target in red. The video demonstrates microrobots attempting to follow the path in real time.

**Movie S4.** Transfer learning from a simulation environment to a real vascular channel using an MBRL model with sweeping actions.

**Movie S5.** MBRL general model trained on 10 environments demonstrates its ability to perform across all 10 environments and adapt to a new, unseen channel with just 30 minutes of additional training.

**Movie S6.** Autonomous manipulation in a flow environment after transfer learning from a simulation that mimics the flow, guiding the microrobot to move in a low-drag region near the wall.
